# Extrinsic Apoptosis and Necroptosis in Telencephalic Development: A Single-Cell Mass Cytometry Study

**DOI:** 10.1101/2025.03.01.640843

**Authors:** Jiachen Shi, Weile Liu, Alison Song, Timi Sanni, Amy Van Deusen, Eli R. Zunder, Christopher D. Deppmann

## Abstract

Regulated cell death is integral to sculpting the developing brain, yet the relative contributions of extrinsic apoptosis and necroptosis remain unclear. Here, we leverage single-cell mass cytometry (CyTOF) to characterize the cellular landscape of the mouse telencephalon in wild-type (WT), RIPK3 knockout (RIPK3 KO), and RIPK3/Caspase-8 double knockout (DKO) mice. Strikingly, combined deletion of RIPK3 and Caspase-8 leads to a 12.6% increase in total cell count, challenging the prevailing notion that intrinsic apoptosis exclusively governs developmental cell elimination. Detailed subpopulation analysis reveals that DKO mice display selective enrichment of Tbr2⁺ intermediate progenitors and endothelial cells, underscoring distinct, cell type–specific roles for extrinsic apoptotic and necroptotic pathways. These findings provide a revised framework for understanding the coordinated regulation of cell number during telencephalic development and suggest potential mechanistic links to neurodevelopmental disorders characterized by aberrant cell death.

## Introduction

During early development, the central nervous system (CNS) generates an excess of neurons and glia, subsequently eliminating up to 50% to ensure proper circuit formation and brain function^1–7^. Regulation of cell number for specific cell types during this process is orchestrated by a combination of factors: cell proliferation, cell differentiation, and regulated cell death. Historically, intrinsic (mitochondrial) apoptosis was considered the primary mechanism driving developmental cell death, a view supported by abundant cleaved Caspase-3 and TUNEL-positive cells^8–10^, as well as by the severe malformations observed in mice lacking key intrinsic apoptotic components such as Bax/Bak, Caspase-9, or Apaf-1^11–14^. However, since both cleaved Caspase-3 and TUNEL positivity can result from death pathways including extrinsic apoptosis, these classical markers alone cannot definitively attribute developmental cell elimination solely to intrinsic apoptosis. Furthermore, even in mice lacking the key intrinsic apoptotic components Apaf-1^-/-^ and Bax^-/-^Bak^-/-^, substantial cell death persists in multiple tissues^15,16^, including interdigital cell death, suggesting the involvement of alternative mechanisms such as extrinsic apoptosis.

To genetically dissect the contribution of extrinsic apoptosis to cell elimination during neural development, the initiator Caspase-8 is an intuitive target because it serves as the convergence point for multiple death receptors such as Fas/CD95, TRAILR1, and TNFR1^17–19^. Upon death receptor activation, procaspase-8 and adaptor protein FADD are recruited to form the death-inducing signaling complex (DISC). DISC assembly promotes procaspase-8 dimerization and autoprocessing to active Caspase-8, which then triggers apoptosis by cleaving and activating executioner Caspase-3^19,20^. Beyond direct caspase activation, Caspase-8 also amplifies the death signal by generating truncated Bid (tBid), which promotes mitochondrial outer membrane permeabilization (MOMP) through both Bax/Bak-dependent and independent mechanisms^21–23^. Given its central role, Caspase-8 would seem to be a straightforward target for studying extrinsic apoptosis in neural development. However, Caspase-8 also negatively regulates other cell death pathways. Consequently, its loss not only disrupts extrinsic apoptosis but also serves as a gain-of-function for alternative death mechanisms, complicating interpretation of the knockout phenotype.

In addition to its canonical role mediating extrinsic apoptosis, Caspase-8 activity also suppresses necroptosis by inhibiting activation of the RIPK1-RIPK3-MLKL signaling cascade^24–28^. The consequence of disrupting this dual function is severe: Caspase-8-deficient mice die embryonically by E11.5 due to unopposed necroptotic signaling, which gives rise to hyperaemia, neural tube undulation and vascular defects/dysfunction^29,30^. Notably, the vascular phenotype appears central to this lethality, as conditional knockout of Caspase-8 in endothelial cells (Casp8^F/−^Tie1-Cre) recapitulates the vascular defects and embryonic lethality of the global knockout, suggesting endothelial cells as a key population vulnerable to necroptotic cell death^31^. The role of necroptosis is further confirmed by the rescue of embryonic lethality through concurrent deletion of RIPK3, demonstrating the critical importance of this regulatory balance during development^27,30^.

Collectively, these findings not only underscore the centrality of extrinsic apoptosis in neural development but also highlight necroptosis as a mechanistically distinct pathway with significant developmental implications. Unchecked activation of the RIPK1-RIPK3-MLKL cascade, as observed in Caspase-8–deficient contexts, suggests that necroptosis contributes directly to vascular homeostasis during embryogenesis. Moreover, the potential for necroptosis to modulate cell numbers—complementing the cell elimination mediated by apoptotic pathways—provides a compelling rationale for its investigation as an additional regulatory mechanism that fine-tunes tissue architecture during telencephalic development.

To systematically investigate the contributions of both extrinsic apoptosis and necroptosis during development, we employed a comparative genetic approach using RIPK3/Caspase-8 double knockout (DKO) mice alongside RIPK3 single knockouts (RIPK3 KO) and wild-type (WT) controls. This experimental design creates a powerful framework to examine both pathways: comparing DKO with RIPK3 KO phenotypes reveals the specific contributions of Caspase-8-mediated extrinsic apoptosis, while comparing RIPK3 KO with wild-type mice illuminates the role of RIPK3-mediated necroptosis. This strategy not only circumvents the necroptosis-mediated embryonic lethality associated with isolated Caspase-8 deletion, but also allows us to examine how these interconnected death pathways operate in parallel during neural development.

To investigate the effect of these knockouts on cell number regulation during neural development, we applied single-cell mass cytometry (CyTOF) measurement to simultaneously quantify multiple protein markers including cleaved Caspase-3 across diverse cell populations^32–34^. We focused on the telencephalon as our model system due to its well-characterized extensive cellular remodeling and neural-vascular interactions during development^35–38^. Here, we leverage CyTOF in conjunction with genetic models (DKO, RIPK3 KO and WT) to dissect how different modes of regulated cell death shape telencephalic development. Our goals are to (1) map the distribution of extrinsic apoptosis and necroptosis across distinct cellular populations; (2) identify compensatory or synergistic interactions among death pathways; and (3) determine how these processes interface with neurovascular development. Through these investigations, we reveal new cell type-specific roles for extrinsic apoptosis and necroptosis and show that the combined loss of RIPK3 and Caspase-8 significantly increases telencephalic cell number—highlighting the previously underappreciated importance of death receptor signaling in neurodevelopment. Our findings establish a framework for understanding how cell death pathways work together to refine brain architecture and may have implications for therapies targeting aberrant cell death in neurodevelopmental disorders^39,40^.

## Results

### Profiling Cell Death and Proliferation in Developing Telencephalon Neural Cell Types

To systematically characterize the temporal dynamics of cellular proliferation and death during mouse brain development, we performed comprehensive re-analysis of single-cell mass cytometry data from our previously published developmental atlas of the CNS with daily timepoints from E11 to P4^33^. To include dying cell populations that were excluded from our previous analysis, we removed the Cisplatin-based viability gate^41^ from our analysis pipeline (Fig. 1A). This approach allowed simultaneous quantification of three distinct cellular states using key markers: Ki67 for proliferation^42,43^, cleaved Caspase-3 (CC3) for apoptosis^8,44^, and Cisplatin for plasma membrane integrity^41^. It is important to note that while Cisplatin is used in clinical settings as a chemotherapeutic agent that enters cells and forms DNA adducts which can trigger apoptosis^41,45^, this process takes place on the hour time scale, and our protocol’s viability stain with Cisplatin only lasts for 30 seconds. This short exposure followed by immediate PFA fixation (Methods) ensures that Cisplatin doesn’t induce apoptosis in our cell samples, instead only acting as a viability dye via elevated staining levels in dying cells with compromised plasma membranes^41^.

**Figure 1.**
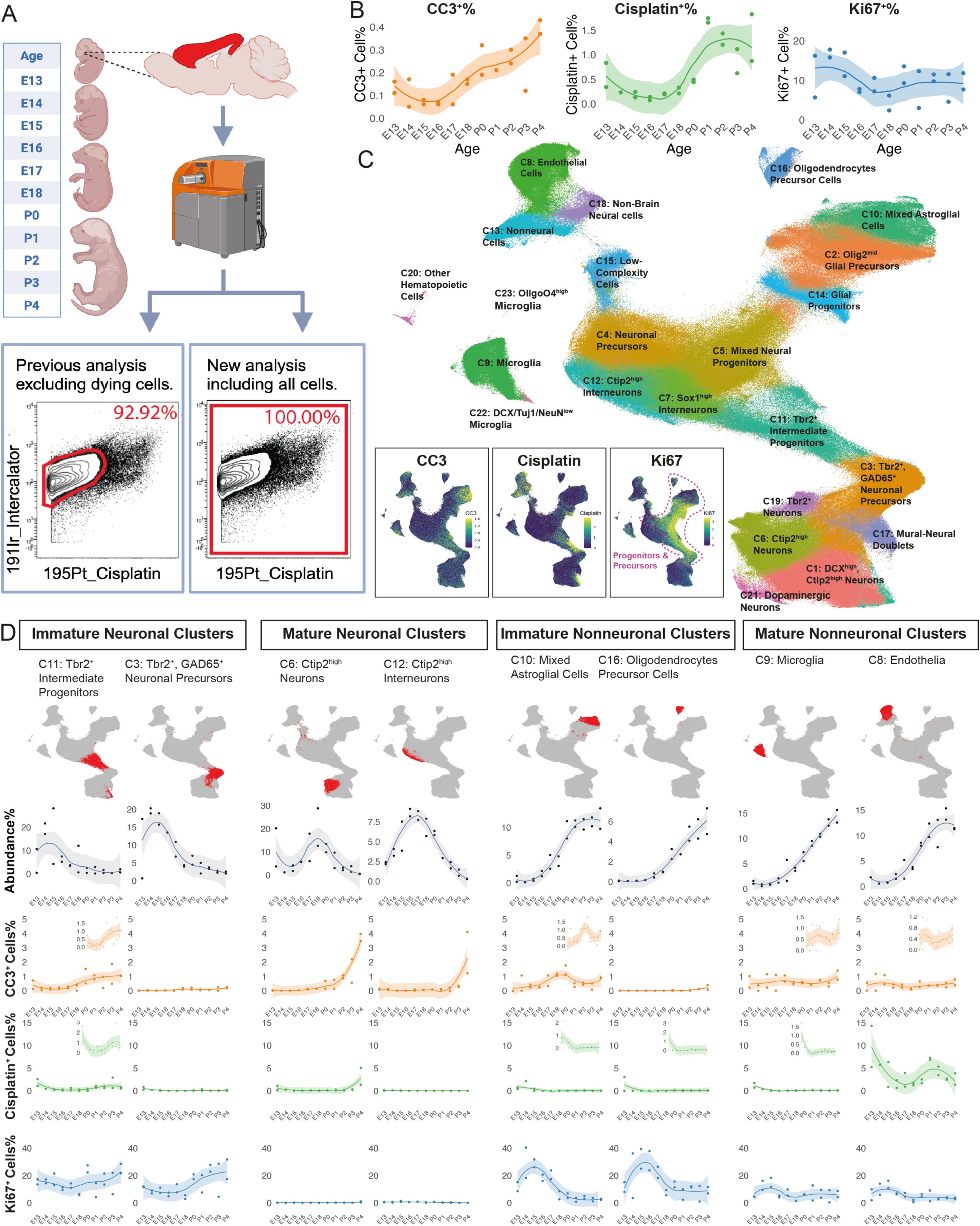
Cell Death, Proliferation, and Abundance in the Developing Telencephalon. (A) Schematic representation of the modified experimental workflow for including both live and dying cells. Single-cell mass cytometry data from mouse telencephalon (E13-P4) were reanalyzed to include both live and dying cells. Flow cytometry plots show the inclusion of Cisplatin^+^ cells in the new analysis. (B) Temporal dynamics of CC3^+^%, Cisplatin^+^%, and Ki67^+^% cells throughout development (E13-P4). Individual replicates are shown along with Loess curve fitting of the data. (C) UMAP visualization of 23 distinct cell clusters identified by Leiden clustering. Insets show the expression patterns of CC3 (apoptosis), Cisplatin (plasma membrane integrity), and Ki67 (proliferation) across the UMAP layout. Cell clusters identified as precursor and progenitor cell types by their protein expression profiles are highlighted by a red dashed line in the Ki67 inset. (D) Detailed analysis of selected clusters representing immature neuronal, mature neuronal, immature nonneuronal, and mature nonneuronal cell populations. Top: UMAP plots highlighting the distribution of each cluster. Bottom: Temporal dynamics of cluster abundance% (gray), CC3^+^ cells% (orange), Cisplatin^+^ cells% (green), and Ki67^+^ cells% (blue) from E13 to P4. Individual replicates are shown along with Loess curve fitting of the data.

Re-analysis including all dead and dying cells in the mouse telencephalon revealed that global cell death progressively increased from E13 to P4, as evidenced by elevated percentages of both CC3^+^ and Cisplatin^+^ cells (Fig. 1B). Between E13 and P4, CC3^+^ cells increased by 203.0% and Cisplatin^+^ cells by 129.9% (R^2^ = 0.72 for CC3^+^ curve fitting and 0.71 for Cisplatin^+^ curve fitting). This temporal pattern aligns with established developmental cell death dynamics, where a major wave of apoptosis occurs during the first postnatal week, eliminating up to 30% of neurons^46,47^. Previous studies have demonstrated that neocortical cell death exhibits characteristic temporal dynamics: initiating at P2, reaching maximal levels during P4-P7, followed by gradual reduction through P9-P10^48^.

In addition to these temporal changes, we identified distinct cell populations exhibiting specific combinations of death markers (Supplementary Fig. S1A): CC3^+^Cisplatin^-^ cells, CC3^-^Cisplatin^+^ cells, and CC3^+^Cisplatin^+^ cells. Based on established cell death mechanisms, where classical apoptosis typically maintains membrane integrity while non-apoptotic pathways such as necroptosis are characterized by early membrane rupture^49^, these populations were interpreted as follows: CC3^+^Cisplatin^-^ cells represent early apoptotic cells with intact membrane integrity; CC3^-^Cisplatin^+^ events suggest cells undergoing non-apoptotic death with compromised membranes; and CC3^+^Cisplatin^+^ cells potentially indicate a combination of later-stage apoptotic and non-apoptotic death^49,50^. This heterogeneity in death marker profiles hints at an interplay between different cell death mechanisms during development.

While cell death showed temporal progression, we did not observe a clear pattern in proliferation (Fig. 1B). The percentage of Ki67^+^ ranges from 2.35% to 17.77% across developmental stages, without a simple trend (R² = 0.31 for Ki67^+^ curve fitting). This complex pattern at the population level likely reflects the partially overlapping waves of neurogenesis (E12 through late embryonic stage) and gliogenesis (late embryonic stage through early postnatal stage)^51–53^, during which proliferative activity transitions between different progenitor populations. To uncover cell type - specific patterns that might be masked by these population averages, we next performed detailed analyses of distinct cell populations.

To characterize distinct cell populations, we performed Leiden clustering^54^ and visualized the results using Uniform Manifold Approximation and Projection (UMAP)^55^. The analysis revealed 23 distinct clusters (Fig. 1C, Supplementary Fig. S1B), which were subsequently grouped into four broad categories based on their unique protein expression profiles: 1) immature neuronal clusters, 2) mature neuronal clusters, 3) immature nonneuronal clusters, and 4) mature nonneuronal clusters. This classification was established using differential expression patterns of validated markers for neuronal maturity (e.g., PAX6, Tbr2, DCX, Tbr1, Ctip2)^56–58^ and glial cell types (e.g., GFAP, Olig2, CD11b, PECAM)^59–61^.

Analysis of proliferation and death markers across the UMAP layout revealed distinct maturation-associated patterns (Fig. 1C insets). Ki67 exhibited high intensity in progenitor and precursor clusters, particularly in mixed neural progenitors, glial progenitors, and Tbr2^+^ intermediate progenitors. This enrichment of the proliferation marker in progenitor and precursor populations aligns with established patterns of active cell division in neural progenitor cells during CNS development^62^. Conversely, CC3 and Cisplatin showed higher intensity in more differentiated populations, notably in mixed astroglial cells and endothelial cells.

Detailed temporal analysis of individual clusters revealed distinct death and proliferation patterns in neuronal populations (Fig. 1D, Supplementary Fig. S1C). Most immature neuronal clusters maintained high percentages of Ki67^+^ cells (>10%) while showing low percentages of both CC3^+^ and Cisplatin^+^ cells (0-2%). In contrast, mature neuronal clusters generally exhibited limited proliferation (Ki67^+^ cells <2%) and low death marker expression (CC3^+^ and Cisplatin^+^ cells 0-2%), although there were some notable exceptions. Specifically, Ctip2^high^ neurons and Ctip2^high^ interneurons showed progressive increases in apoptotic cells (CC3^+^% increasing from 0% at E13.5 to around 4% at P4). While mature neurons generally exhibit higher resistance to death signals in the adult brain^63^, our analysis window (E13 to P4) includes a critical period of developmental apoptosis (P0-P4) when neurons are particularly susceptible to death signals as circuits are refined and excess cells are eliminated^46,47,64^.

Nonneuronal clusters exhibited more complex patterns (Fig. 1D, Supplementary Fig. S2). The relative abundance of most nonneuronal clusters started to increase around E15-E18 with astrogenesis starting around E16-E17. This timing aligns with the established developmental program where gliogenesis follows neurogenesis, with astrocyte production beginning in late embryonic stages and continuing through early postnatal development^51^. While immature nonneuronal clusters generally maintained high percentages of Ki67^+^ cells (>15%), some mature nonneuronal clusters also displayed significant Ki67^+^ expression (5-20%), consistent with the established pattern where glial cells, unlike neurons, retain proliferative capacity throughout development^65,66^. Notably, specific immature nonneuronal clusters (e.g., Olig2^med^ Glial Precursors, Astroglial Precursors, and Oligodendrocyte Precursor Cells) showed a distinct temporal pattern of proliferation, characterized by Ki67^+^% increasing from ∼40% at E13 to a peak of ∼70% around E15-E16, followed by a subsequent decrease to ∼30% by P4.

In terms of cell death dynamics, mature nonneuronal clusters displayed varying levels of death markers. Notably, endothelial cells demonstrated an intriguing discrepancy between markers: while Cisplatin uptake was substantial (5-15% of cells), cleaved Caspase-3 (CC3) expression remained relatively low (<2%). The high Cisplatin positivity coupled with minimal CC3 activation indicates the engagement of alternative cell death pathways such as necroptosis in developmental vascular pruning. Though CC3 analysis captured both intrinsic and extrinsic apoptotic pathways, and Cisplatin indicated potential necroptotic death, the unexpected divergence between these markers prompted us to investigate specific death pathways through genetic perturbation of extrinsic apoptosis and necroptosis.

### Molecular Architecture of Death Receptor Signaling in the Developing Telencephalon

To dissect the molecular mechanisms underlying the observed cell death patterns across embryonic and postnatal development in the mouse telencephalon, we examined the complex interplay between death receptor-mediated pathways and mitochondrial apoptosis (Fig. 2A). In wild-type conditions, RIPK1 can either engage FADD/Caspase-8-dependent apoptotic pathway or trigger RIPK3/MLKL-mediated necroptosis^27,28^. Once activated, Caspase-8 not only directly promotes apoptosis by Caspase-3 cleavage and activation, but also engages mitochondrial amplification through cleavage of Bid to generate truncated Bid (tBid), which translocates to mitochondria and induces MOMP through both Bax/Bak-dependent and independent mechanisms, though the latter operates with reduced efficiency^21–23^. This mitochondrial loop is especially crucial in Type II cells, where direct Caspase-8 signaling alone may be insufficient to execute the death program^67^. Genetic ablation of RIPK3 (RIPK3 KO) eliminates the necroptotic branch while maintaining both direct and mitochondria-amplified apoptotic potential. Conversely, Caspase-8 deficiency proves embryonically lethal due to unopposed necroptosis^29,30^. The combined deletion of RIPK3 and Caspase-8 (DKO) functionally disables both pathways, providing a unique model to investigate their coordinated roles in developmental cell death.

**Figure 2.**
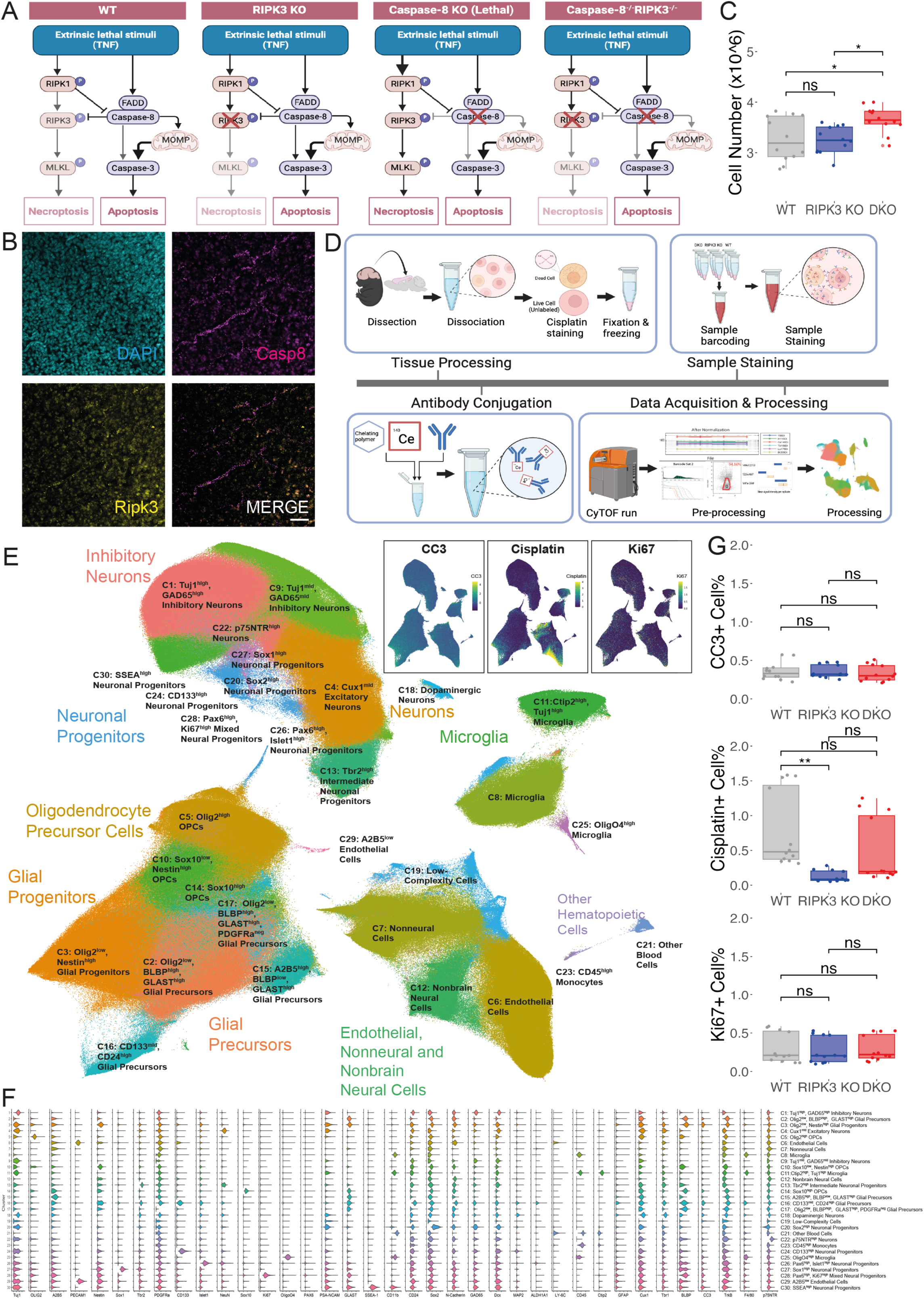
Investigation of Extrinsic Apoptosis and Necroptosis in Telencephalon Development. (A) Schematic representation of extrinsic apoptotic and necroptotic pathways in WT, RIPK3 KO, Caspase-8 KO, and DKO mice. (B) Representative images of RNAscope in situ hybridization for Casp8 (magenta) and Ripk3 (yellow) mRNA in P4 WT mouse cortex. DAPI (cyan) labels nuclei. Scale bar: 50 μm. (C) Quantification of total cell numbers in the telencephalon of P4 mice (n = 12 per genotype). (D) Workflow schematic for mass cytometry analysis of P4 mouse telencephalon from WT, RIPK3 KO and DKO mice, including tissue processing, antibody conjugation, sample staining, and data acquisition and processing steps. (E) UMAP visualization of mass cytometry data showing 30 distinct cell clusters identified by Leiden clustering. Major cell type categories are annotated. Insets show expression patterns of CC3, Cisplatin, and Ki67 across the UMAP layout. (F) Violin plot showing relative expression of all 37 markers across the 30 identified cell clusters. (G) Quantification of CC3^+^, Cisplatin^+^, and Ki67^+^ cells% across genotypes from mass cytometry data. For panels C and G, data presented as box plots where the middle line represents the median, the box represents the interquartile range (IQR), and the whiskers extend to 1.5 times the IQR. Individual data points are shown as dots. Statistical differences between groups are denoted by asterisks. Statistical significance levels are as follows: *p<0.05,* * p<0.01, *** p<0.001, ns: not significant (one-way ANOVA with Tukey’s post-hoc test).

### Expression of Extrinsic Apoptosis and Necroptosis Components in the Telencephalon

We initially attempted to detect pathway-specific proteins - cleaved Caspase-8 (CC8) for extrinsic apoptosis and phosphorylated MLKL (pMLKL) for necroptosis - via immunohistochemistry (IHC). However, extensive validation testing of multiple commercial antibodies revealed lack of specificity, as demonstrated by inconsistent staining patterns and high background signal (Supplementary Fig. S3 and S4). Given the strong developmental regulation of death receptors and their ligands, with peak expression at P5, and minimal detection in adult brain^68^, we hypothesized that analyzing mRNA expression patterns might reveal regulatory mechanisms governing apoptotic and necroptotic pathways. Using RNAscope to examine P4 WT mouse cortex (n = 3), we observed widespread Casp8 mRNA expression, with notable colocalization of Casp8 and RIPK3 mRNA in cells with endothelial morphology (Fig. 2B). This pattern was specific to developing brains, as no similar colocalization was observed in adult cortex (Supplementary Fig. S5). The co-expression of these death pathway components in endothelial cells is particularly intriguing given that endothelial disorganization and subsequent circulatory failure has been identified as the primary cause of embryonic lethality in Caspase-8 knockout mice^30,31^. This spatial convergence suggests potential crosstalk between these death mechanisms in endothelial cells.

To assess the functional significance of these pathways, we extended our study to include RIPK3 KO and DKO mice alongside WT controls. These mouse models have been previously validated for studying the roles of necroptosis and extrinsic apoptosis in development^30,69^. We focused our investigation on P4 telencephalon for two strategic reasons: first, this represents a critical window to capture the cumulative effects of both late embryonic and early postnatal waves of cell death^46,48^; second, this time point allows optimal preparation of high-quality single-cell suspensions for downstream CyTOF analysis^33,70^.

### Impact of Death Receptor Pathway Deletions on Telencephalic Cell Number

Initial phenotypic characterization using automated cell counting of telencephalon revealed no significant difference in total cell count between RIPK3 KO (3.25 ± 0.27 × 10^6^ cells) and WT groups (3.27 ± 0.44 × 10^6^ cells; p = 0.986). However, the DKO group exhibited a significantly higher cell count (3.66 ± 0.26 × 10^6^ cells) compared to both WT and RIPK3 KO groups (p < 0.05 for both comparisons) (Fig. 2C). This effect was consistent across both sexes (Supplementary Fig. S6), with analysis of three independent litters per genotype, using two males and two females per litter (n = 12 per genotype). The significant increase in cell number specifically in DKO mice (12-13% higher than both WT and RIPK3 KO, p < 0.05), coupled with the lack of effect in RIPK3 single knockouts, that Caspase-8-mediated extrinsic apoptosis significantly contributes to cell elimination during telencephalon development. While we cannot completely exclude synergistic effects or compensatory mechanisms triggered by combined pathway deletion, these findings establish death receptor signaling as a key regulator of developmental cell death.

To elucidate the cellular mechanisms underlying the observed increases in cell number, we performed high-dimensional mass cytometry using a 37-antibody panel (Fig. 2D, Supplementary Table 1) on 3,604,163 single cells from the developing telencephalon. Initial Leiden clustering identified 30 distinct clusters (Fig. 2E and 2F), with UMAP visualization showing clear segregation of major cell populations based on established markers^33^ (Supplementary Fig. S7): neurons (NeuN^+^, MAP2^+^, TuJ1^+^), astrocytes (GFAP^+^, ALDH1A1^+^), oligodendrocytes (Olig2^+^, OligO4^+^, Sox10^+^), microglia (CD45^+^, CD11b^+^), and endothelial cells (CD31^+^, Ly6C^+^). To achieve higher resolution, we performed subclustering analysis on four major cellular categories: Neuronal populations; Glial precursor populations; Endothelial, Astrocytic, and Nonneural populations; and Microglial, Blood, and Monotype populations. This refined analysis revealed 89 distinct subpopulations (Supplementary Fig. S8, 9).

Analysis of cell death markers revealed three key findings (Fig 2G): **1)** Classic apoptotic markers remained unchanged across genotypes (CC3^+^ cells: WT 0.358 ± 0.116%, RIPK3 KO 0.355 ± 0.084%, DKO 0.329 ± 0.106%; p > 0.05). **2)** Proliferation rates showed no significant differences (Ki67^+^ cells: WT 0.298 ± 0.195%, RIPK3 KO 0.272 ± 0.170%, DKO 0.293 ± 0.166%; p > 0.05). **3)** Membrane permeability showed striking genotype-dependent effects with RIPK3 KO mice exhibiting significantly reduced Cisplatin uptake compared to WT (0.125 ± 0.082% vs 0.788 ± 0.552%, p < 0.01) and DKO mice showing intermediate levels (0.481 ± 0.481%), suggesting compensatory mechanisms that partially restore membrane permeability^71^. While this global analysis reveals intriguing population-level changes across genotypes, definitive determination of cell type-specific roles for these death pathways requires higher resolution analysis of each cellular compartment.

To explore cell type-specific changes, we started with hierarchical clustering analysis of marker expression across 30 initial clusters, revealing distinct molecular signatures associated with genotype (Supplementary Fig. S10). The resulting heatmap demonstrates clear segregation of cellular clusters based on lineage-specific markers. Cross-genotype comparisons revealed significant population changes, particularly between DKO vs. WT and RIPK3 KO vs. WT, as shown in the p-value heatmap (Supplementary Fig. S9, right panels). These changes were especially pronounced in glial and non-neuronal populations, including microglia (C8, C25), endothelial cells (C6), low-complexity cells (C19), nonbrain cells (C12), and blood cells (C21), suggesting cell type-specific regulation by extrinsic apoptosis and necroptosis.

### Selective Expansion or Depletion of Neuronal Populations Following Death Receptor Pathway Disruption

Given these broad population changes, we next performed detailed characterization of specific cellular compartments. Comprehensive analysis of neuronal populations across genotypes revealed distinct alterations in both progenitor and mature neuronal compartments, with particularly striking differences in intermediate progenitor populations. Initial clustering analysis identified four key neuronal populations with genotype-specific abundance patterns (Fig. 3A): Tbr2^high^ Intermediate Neuronal Progenitors (C13), Cux1^mid^ Excitatory Neurons (C4), Dopaminergic Neurons (C18), and p75NTR^high^ Neurons (C22). Among these, DKO mice exhibited a selective increase in Tbr2^high^ Intermediate Neuronal Progenitors (2.63 ± 0.22%) compared to both WT (2.30 ± 0.36%, p < 0.05) and RIPK3 KO (2.45 ± 0.27%) populations.

**Figure 3.**
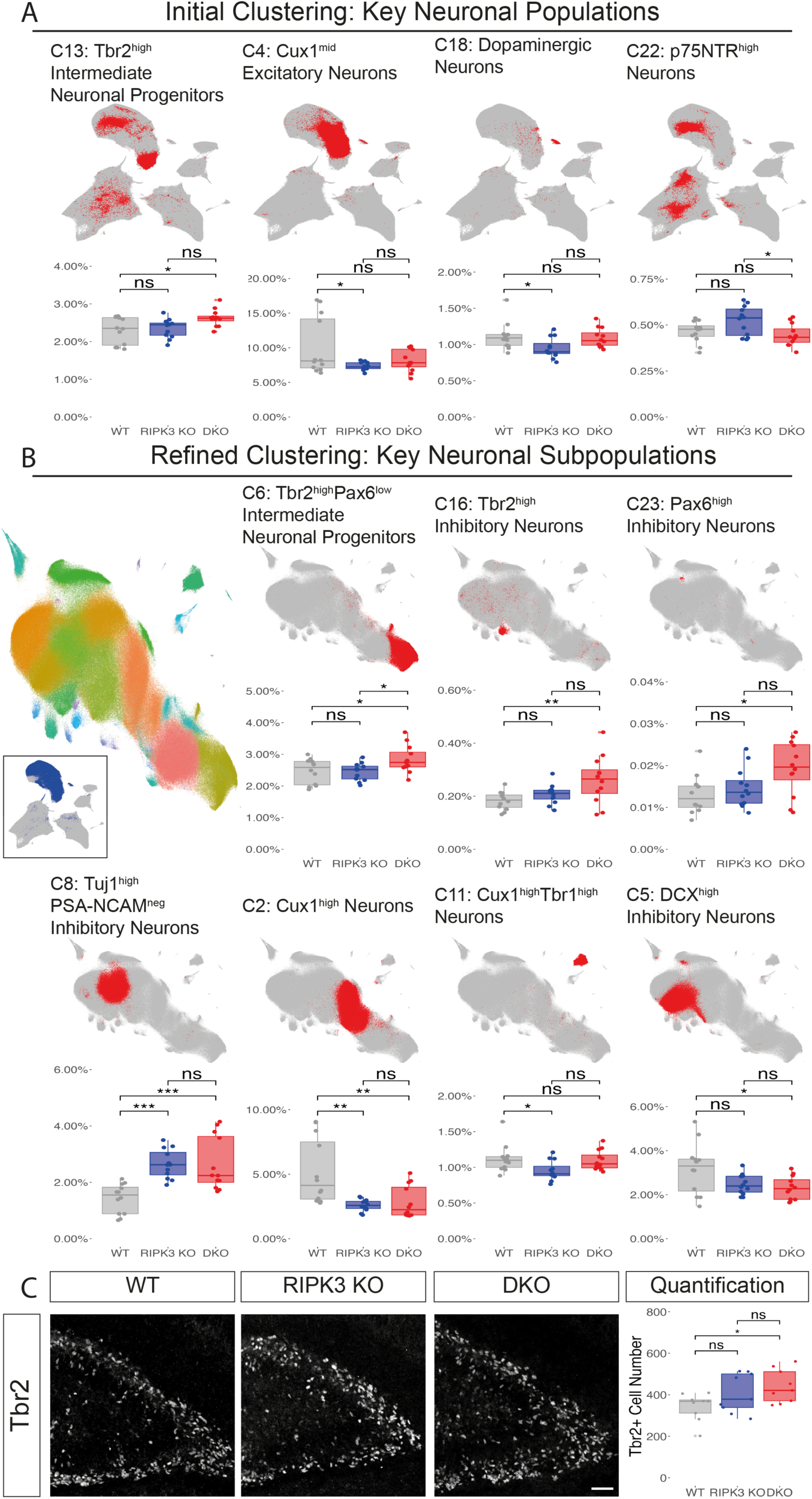
Differential Abundance of Telencephalic Neuronal Subpopulations. (A) Initial clustering analysis showing UMAP visualization and quantification of four major neuronal populations. For each cluster, relative abundance% is compared across WT, RIPK3 KO, and DKO. (B) Refined clustering analysis showing UMAP visualization and quantification of key neuronal subpopulations. Left UMAP plot displays 26 distinct neuronal subclusters (C1-C26) identified through iterative Leiden clustering (colored overlay), with inset showing the analyzed clusters from initial clustering (dark blue). Representative UMAP plots and quantification are shown for seven key neuronal populations. For each cluster, relative abundance% is compared across WT, RIPK3 KO, and DKO. (C) IHC validation of Tbr2^+^ cells. Left: Representative images of Tbr2 staining in the dentate gyrus across genotypes. Right: Quantification of Tbr2^+^ cell numbers in the dentate gyrus. Scale bar: 50 μm. For all quantifications: Data are presented as box plots showing median (middle line), interquartile range (box), and whiskers (1.5×IQR), with individual data points overlaid. Statistical significance determined by one-way ANOVA with Tukey’s post-hoc test (*p<0.05, **p<0.01, ***p<0.001, ns: not significant).

Further subclustering of neuronal populations revealed more nuanced genotype-dependent differences (Fig. 3B). Tbr2^high^Pax6^low^ Intermediate Neuronal Progenitors (C6) showed significant enrichment in DKO mice (2.84 ± 0.43%) compared to both WT (2.46 ± 0.39%, p < 0.05) and RIPK3 KO (2.45 ± 0.27%, p < 0.05). Similarly, Tbr2^high^ Inhibitory Neuron population (C16) exhibited increased abundance in DKO mice (0.26 ± 0.09%) versus WT controls (0.18 ± 0.03%, p < 0.01). We also observed elevated proportions of Pax6^high^ Inhibitory Neurons (C23) in DKO mice (0.019 ± 0.0065%) compared to WT controls (0.013 ± 0.0046%, p < 0.05). Investigation of mature neuronal populations revealed additional genotype-specific differences. Both RIPK3 KO (2.65 ± 0.51%) and DKO (2.66 ± 0.95%) mice showed significant increases in Tuj1^high^PSA-NCAM^neg^ Inhibitory Neurons (C8) compared to WT controls (1.43 ± 0.52%, p < 0.001). Conversely, Cux1^high^ neurons (C2) decreased in both RIPK3 KO (2.60 ± 0.51%) and DKO mice (2.85 ± 0.51%) compared to WT controls (5.17 ± 2.52%, p < 0.01). Additionally, Cux1^high^Tbr1^high^ Neurons (C11) were significantly reduced in RIPK3 KO mice (0.95 ± 0.14%) compared to WT (1.12 ± 0.19%, p < 0.05), while DKO mice exhibited decreased DCX^high^ inhibitory neurons (C5) (1.12 ± 0.31% vs WT: 1.85 ± 0.42%, p < 0.05).

To validate the mass cytometry findings, we performed IHC analysis of Tbr2^+^ cells in the dentate gyrus (Fig. 3C). This orthogonal approach confirmed increased Tbr2^+^ cell numbers in DKO mice (441 ± 38 cells) compared to WT controls (343 ± 29 cells, p < 0.05), with RIPK3 KO mice displaying an intermediate phenotype (372 ± 33 cells) not significantly different from either WT or DKO (p > 0.05).

### Caspase-8 is Required for Restricting Neurovascular and OPC Cell Number

Analysis of nonneuronal populations revealed striking genotype-dependent alterations in cellular composition, with particularly dramatic effects on vascular and glial lineages. Initial clustering analysis identified four distinct nonneuronal populations exhibiting genotype-specific abundance patterns (Fig. 4A): Endothelial Cells (C6), Sox10^high^ OPCs (C14), OligO4^high^ microglia (C25), and Nonbrain Neural Cells (C12). The most pronounced differences emerged in the endothelial compartment, where DKO mice displayed significantly elevated populations (8.45 ± 0.83%) compared to WT controls (6.95 ± 0.96%, p < 0.01). (Fig. 4A). This vascular expansion was accompanied by marked changes in oligodendrocyte lineage cells, with Sox10^high^ OPCs showing significant enrichment in DKO mice (2.06 ± 0.30%) relative to both WT (1.58 ± 0.65%, p < 0.05) and RIPK3 KO (1.40 ± 0.37%, p < 0.01) groups (Fig. 4A). The selective expansion of Sox10 ^high^ populations suggests altered oligodendrocyte specification and maturation dynamics, consistent with Sox10’s established role as a key regulator of oligodendrocyte development^72^. We also observed a higher percentage of OligO4^high^ microglia (C25) in RIPK3 KO mice (0.20 ± 0.02%) compared to both WT (0.16 ± 0.03%, p < 0.01) and DKO (0.14 ± 0.02%, p < 0.001) groups. The co-expression of OligO4 with canonical microglial markers (CD11b and CD45) suggests these cells may represent a specialized population engaged in myelin phagocytosis, aligning with previously characterized myelin-engulfing microglia population^73–75^.

**Figure 4.**
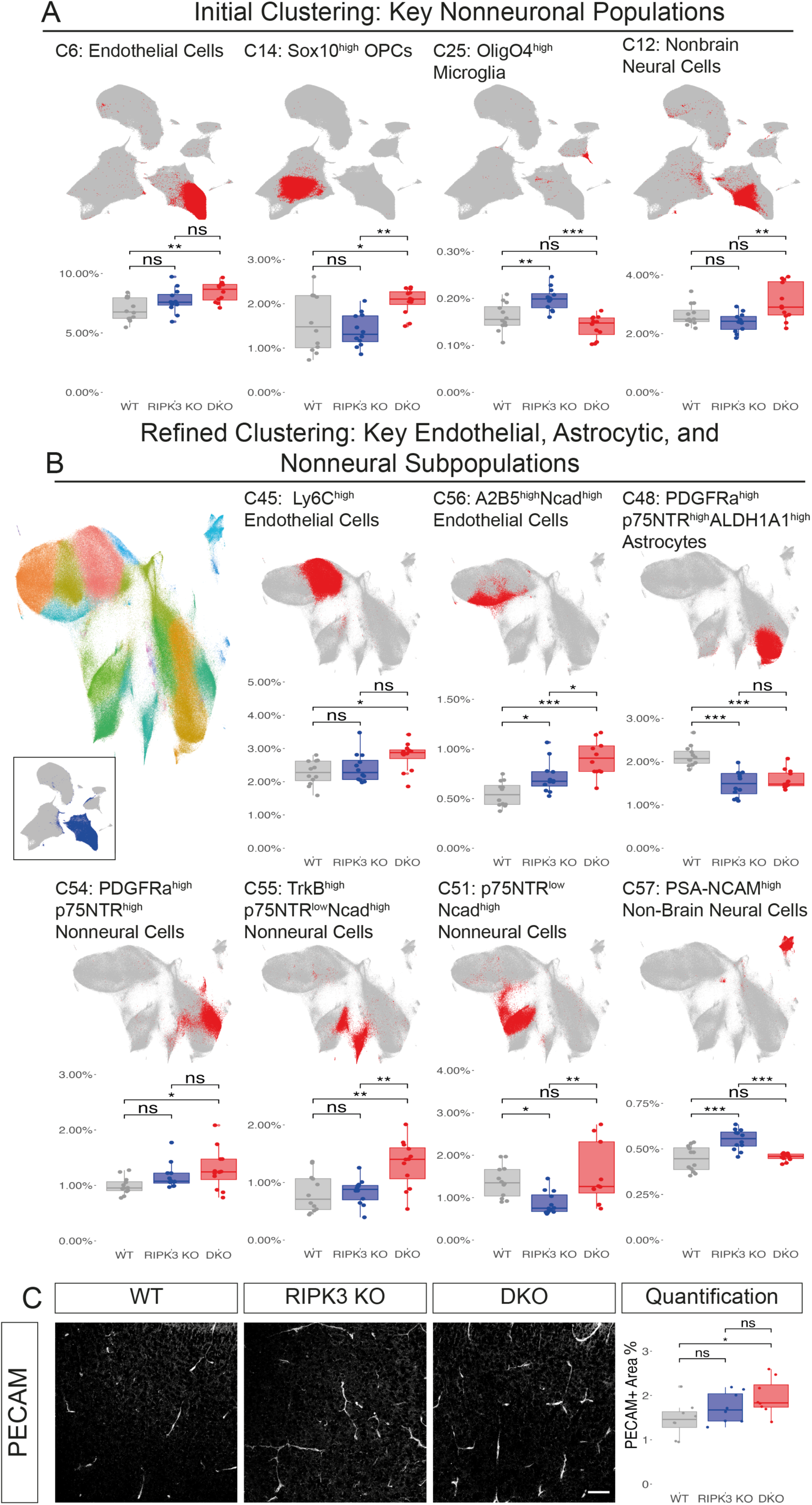
Differential Abundance of Telencephalic Nonneuronal Subpopulations. (A) Initial clustering analysis showing UMAP visualization and quantification of four major nonneuronal populations. For each cluster, relative abundance% is compared across WT, RIPK3 KO, and DKO. (B) Refined clustering analysis showing UMAP visualization and quantification of key endothelial, astrocytic, and nonneural subpopulations. Left UMAP plot displays 22 distinct nonneuronal subclusters (C45-C66) identified through iterative Leiden clustering (colored overlay), with inset showing the analyzed clusters from initial clustering (dark blue). Representative UMAP plots and quantification are shown for seven key endothelial, astrocytic, and nonneural subpopulations. For each cluster, relative abundance% is compared across WT, RIPK3 KO, and DKO. C) IHC validation of PECAM+ area. Left: Representative images of PECAM staining in the cortex across genotypes. Right: Quantification of PECAM+ Area% in the cortex. Scale bar: 50 μm.

Through secondary clustering, we identified distinct endothelial subpopulations with differential abundance across genotypes. Ly6C^high^ Endothelial Cells (C45) were significantly enriched in DKO mice (2.77 ± 0.42%) compared to WT (2.27 ± 0.37%, p < 0.05). Additionally, A2B5^high^Ncad^high^ Endothelial Cells (C56) showed significant elevation in both RIPK3 KO (0.72 ± 0.16%, p < 0.05) and DKO mice (0.91 ± 0.17%, p < 0.001) compared to WT controls (0.55 ± 0.12%), and further enrichment in DKO compared to RIPK3 KO mice (p < 0.05). We also observed pronounced differences in nonneural populations. DKO showed significant increase in both PDGFRa^high^p75NTR^high^ Nonneural Cells (C54) (1.29 ± 0.37% vs WT: 0.99 ± 0.15%, p < 0.05) and TrkB^high^p75NTR^low^Ncad^high^ Nonneural Cells (C55) (1.33 ± 0.42% vs WT: 0.81 ± 0.34%, p < 0.01). Additionally, p75NTR^low^Ncad^high^ Nonneural Cells (C51) showed a distinct pattern, with RIPK3 KO (0.87 ± 0.27%) showed significant lower abundance compared to both WT (1.38 ± 0.38%, p < 0.05) and DKO (1.56 ± 0.72%, p < 0.01). These populations likely represent vascular smooth muscle cells based on their marker profiles^76–78^. The concurrent expansion of these populations with endothelial cells suggests coordinated regulation of the neurovascular unit during development. We also noticed a significant decrease of PDGFRa^high^p75NTR^high^ALDH1A1^high^ Astrocytes (C48) in both RIPK3 KO (1.49 ± 0.29%) and DKO (1.58 ± 0.22%), compared with WT (2.13 ± 0.24%).

To validate these mass cytometry findings, we performed PECAM (CD31) immunostaining to assess vascular coverage. We found increased vascular coverage in the DKO cortex (1.97 ± 0.15%) compared to WT controls (1.47 ± 0.11%, p < 0.05), with RIPK3 KO mice displayed an intermediate phenotype (1.62 ± 0.13%) (Fig. 3C).

### Extrinsic Apoptosis and Necroptosis are Differentially Required to Regulate Select Glia Precursor Numbers

Detailed subclustering analysis of glial precursor populations revealed distinct effects of RIPK3 and Caspase-8 deletion on specific cellular subsets (Fig. 5A). Within the glial lineage, we identified three key precursor populations with differential responses to genetic perturbation: Olig2^low^BLBP^high^ Glial Precursors (C27), Sox10^high^ Oligodendrocyte Precursors (C35), and Olig2^low^BLBP^neg^ Glial Precursors (C36). Notably, RIPK3 deletion triggered opposing effects in BLBP-expressing versus BLBP-negative populations: C27 (BLBP^high^) cells expanded significantly in RIPK3 KO mice (5.63 ± 1.48% vs WT: 4.33 ± 1.00%, p < 0.05), while C36 (BLBP^neg^) cells showed marked reduction (1.66 ± 0.38% vs WT: 2.18 ± 0.64%, p < 0.05). This divergent response suggests cell type-specific roles for death receptor signaling in regulating distinct glial precursor populations.

**Figure 5.**
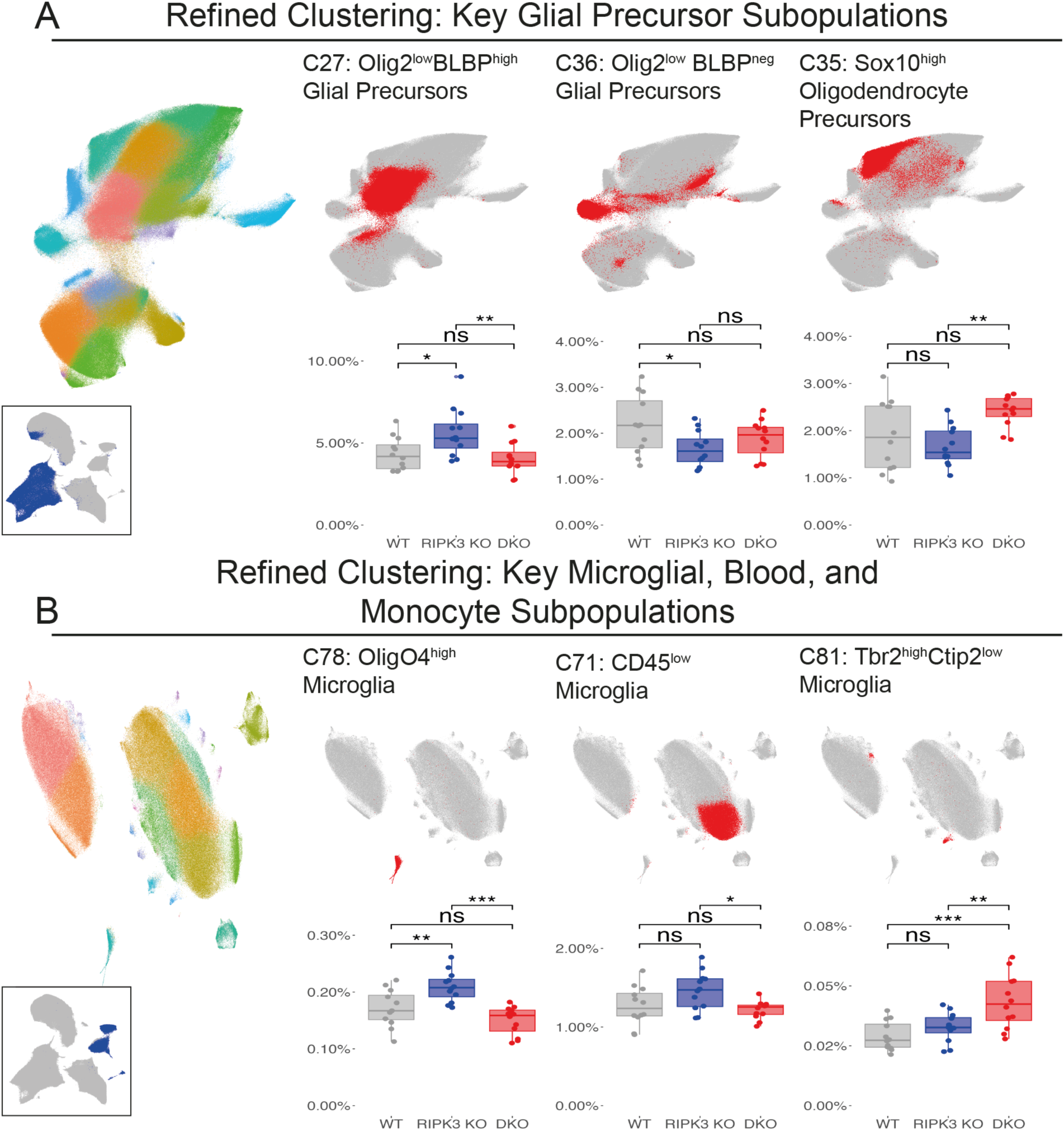
Subclustering Analysis Reveals Differential Glial Subpopulation Responses to Necroptotic and Extrinsic Apoptotic Pathway Disruption. (A) Refined clustering analysis showing UMAP visualization and quantification of key glial lineage subpopulations. Left UMAP plot displays 22 distinct glial subclusters (C27-C44) identified through iterative Leiden clustering (colored overlay), with inset showing the analyzed clusters from initial clustering (dark blue). Representative UMAP plots and quantification are shown for three key glial subpopulations. (B) Refined clustering analysis showing UMAP visualization and quantification of key microglial, blood, and monocyte subpopulations. Left UMAP plot displays 23 distinct microglial, blood, and monocyte subclusters (C67-C89) identified through iterative Leiden clustering (colored overlay), with inset showing the analyzed clusters from initial clustering (dark blue). Representative UMAP plots and quantification are shown for three microglial, blood, and monocyte subpopulations. For each cluster, relative abundance% is compared across WT, RIPK3 KO, and DKO.

### Differential Effects of Death Receptor Pathway Disruption on Cargo Laden Versus Unladen Microglia Numbers

Reclustering of microglial, blood and monocyte subpopulations, we identified three key microglial subpopulations with differential responses to genetic perturbation: CD45^low^ Microglia (C71), OligO4^high^ Microglia (C78), and Tbr2^high^Ctip2^low^ Microglia (C81) (Fig. 5B). The OligO4^high^ subpopulation showed particular sensitivity to RIPK3 deletion, with significant expansion in RIPK3 KO mice (0.21 ± 0.03%) compared to both WT (0.17 ± 0.03%, p < 0.01) and DKO groups (0.15 ± 0.02%, p < 0.001). This finding suggests a specific role for RIPK3-dependent signaling in regulating this microglial subset, independent of Caspase-8 function.

## Discussion

Our single-cell mass cytometry analysis of the developing telencephalon reveals that extrinsic apoptosis and necroptosis operate in parallel with intrinsic apoptosis to regulate cell number during CNS development. In our study, the combined deletion of RIPK3 and Caspase-8 resulted in a 29% expansion of Tbr2⁺ intermediate progenitors and a 34% increase in vascular coverage, while cleaved Caspase-3 (CC3) levels remained largely unchanged. These quantitative changes challenge the traditional view that intrinsic apoptosis alone governs developmental cell death^8–14^ and instead underscore the nonredundant, cell type–specific functions of both death receptor-mediated pathways^18,19,24–26,79,80^.

Cell numbers in development reflect a dynamic equilibrium between proliferation, differentiation, and death (Fig. 6A). The observed changes in population percentages may arise through multiple mechanisms, including intrinsic apoptosis, extrinsic apoptosis, necroptosis, cell state transitions, and compensatory responses. The contribution of each death pathway varies by cell type, as evidenced by the population-specific responses to RIPK3 and Caspase-8 deletion we observed.

**Figure 6.**
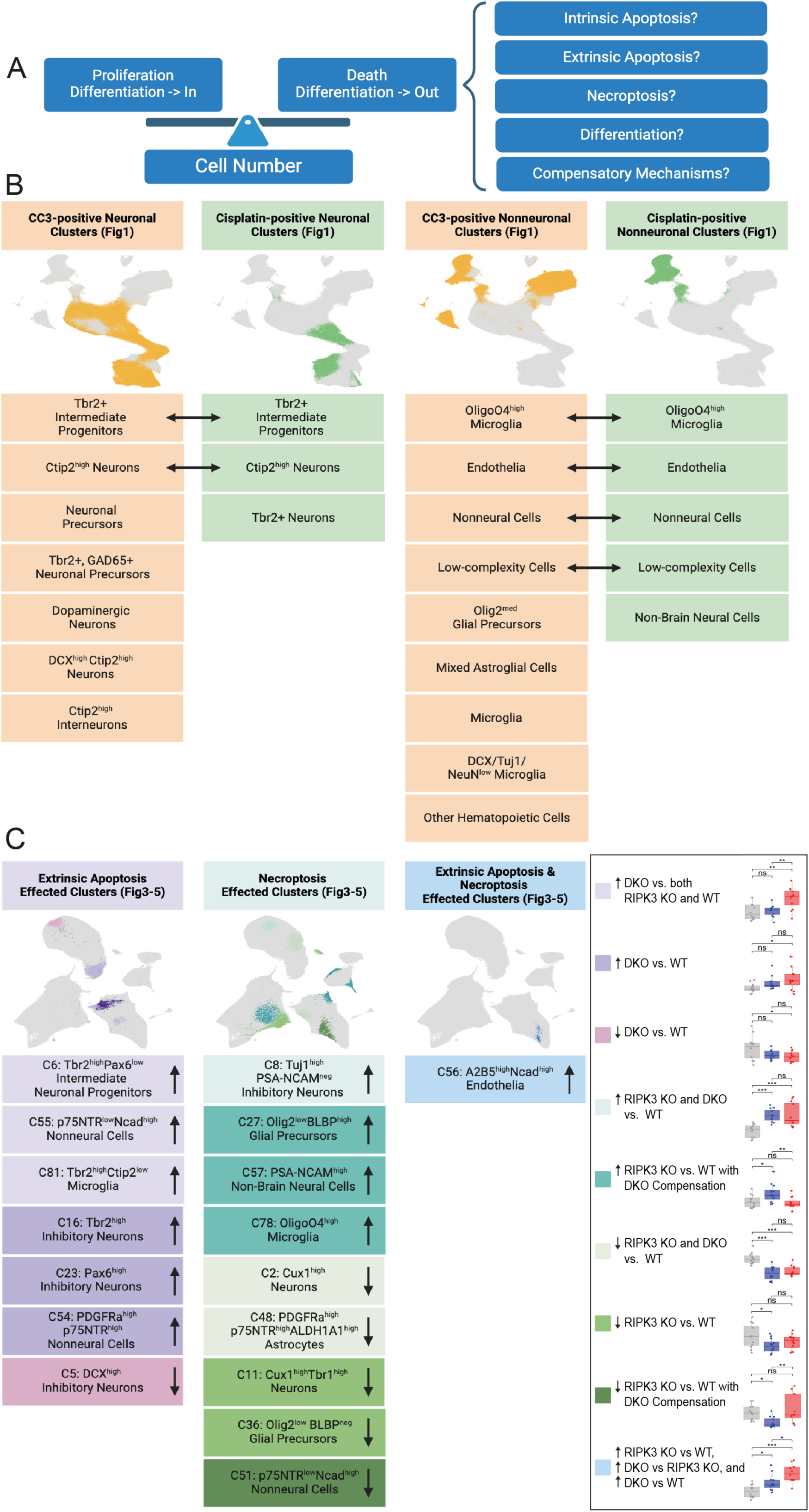
Comprehensive Analysis of Death Pathway-Specific Effects on Telencephalic Cell Population Dynamics. (A) Model of developmental cell number regulation. Left: Cell numbers are balanced between input processes (proliferation and differentiation into the population) and output mechanisms (cell death and differentiation into other populations). Right: Potential mechanisms regulating developmental cell reduction, including distinct death pathways (apoptosis, necroptosis), lineage transitions, and compensatory responses. (B) Summary of cell populations with distinct death signatures in the developing telencephalon (Fig. 1, Supplementary Fig. S1, S2). UMAP visualization (top) and cluster identities (bottom) for neuronal (left) and nonneuronal (right) populations. Clusters showing cleaved Caspase-3 positivity are highlighted in orange, while those with Cisplatin uptake are shown in green. Bidirectional arrows indicate populations positive for both markers. (C) Summary of cell populations affected by death pathway deletion (Fig. 3-5). UMAP visualization (top) and cluster identities (bottom) showing populations regulated by: extrinsic apoptosis (left, purple, unique to DKO), necroptosis (middle, teal/green, RIPK3-dependent), or both pathways (right, blue, sequential changes across genotypes). Arrows indicate increased (↑) or decreased (↓) cluster abundance in knockout conditions.

This cell type-specific regulation is clearly illustrated in our reanalysis of E13-P4 WT telencephalon (Fig. 6B), where we observed distinct death signatures across neuronal and non-neuronal populations. Several populations show evidence of utilizing multiple death pathways, as indicated by the bidirectional arrows between corresponding cell populations. These dual-positive populations include Tbr2^+^ intermediate progenitors and Ctip2^high^ neurons in the neuronal lineages, as well as OligO4^high^ microglia, endothelia, nonneural cells, and low-complexity cells in the non-neuronal populations. This overlap suggests these populations may employ multiple death mechanisms during development, potentially providing redundancy or context-specific regulation of cell numbers.

Notably, our genetic perturbation studies reveal three distinct patterns of pathway engagement. Some populations are uniquely affected in the RIPK3/Caspase-8 DKO but not in the RIPK3 single knockout, implicating a specific role for extrinsic apoptosis (e.g., Tbr2^high^Pax6^low^ intermediate progenitors). Other populations appear regulated primarily by RIPK3-dependent mechanisms (e.g., Tuj1^high^PSA-NCAM^neg^ inhibitory neurons), whereas certain cell types, such as A2B5^high^Ncad^high^ endothelial cells, require coordinated input from both pathways (Fig. 6C). This diversity in cell death regulation may reflect evolutionary adaptations that tailor death pathway engagement to the developmental requirements of specific cellular contexts^81^.

A particularly striking aspect of our study is the expansion of the endothelial compartment in DKO mice. Our data suggest that telencephalic endothelial cells, which uniquely co-express high levels of Caspase-8 and RIPK3 (Fig. 2B), are primed for pathway switching. The concomitant increase in vascular smooth muscle cells further implies a coordinated regulation of the neurovascular unit, aligning with recent observations that endothelial-derived signals modulate neural progenitor behavior through both contact-dependent and paracrine mechanisms^82,83^. Moreover, the genotype-dependent differences in Cisplatin uptake—where RIPK3 deletion alone significantly reduces membrane permeability (>80% reduction, p < 0.01) but combined deletion partially restores it (61% of WT levels)—suggest the activation of compensatory mechanisms, potentially involving alternative effectors such as gasdermin family proteins or noncanonical caspase cascades^2,28^.

Despite these advances, several key questions remain. Our analyses at a single developmental timepoint (P4) capture a critical window of postnatal pruning^46,48^ but do not resolve the temporal dynamics of pathway engagement. Future studies employing inducible Cre-loxP systems to delete RIPK3 and Caspase-8 at defined developmental stages are needed to delineate stage-specific requirements. In addition, conditional deletion in specific cell types (e.g., endothelial-specific deletion using Tie2-Cre or Cdh5-CreERT2) will help distinguish primary, cell-autonomous effects from secondary consequences on neural populations.

Finally, while our study focuses on extrinsic apoptosis and necroptosis, other regulated cell death mechanisms—such as pyroptosis and ferroptosis—may also contribute to developmental cell death^2,84,85^. Integration of spatial transcriptomics with proteomic profiling of death effector complexes could reveal how these pathways interact to maintain the delicate balance of cell number during brain development, with implications for understanding neurodevelopmental disorders associated with aberrant cell death^86^.

In summary, our findings establish a revised framework in which multiple, context-dependent death pathways collaborate to refine brain architecture. By revealing nonredundant roles for extrinsic apoptosis and necroptosis in regulating specific neural and vascular populations, our study not only challenges longstanding paradigms but also provides new insights into the mechanistic underpinnings of CNS development.

## Materials and Methods

### Animals

All animal husbandry and experiments followed Association for Assessment of Laboratory Animal Care guidelines, and were approved by the University of Virginia Animal Care and Use Committee (protocol no. 3795). Mice were housed under controlled environmental conditions, including a 12-hour light/dark cycle, ad libitum access to food and water, and maintained at 21°C with 45-50% humidity. For experimental purposes, P4 pups were harvested from C57/BL6 wild-type, RIPK3 KO, and DKO mice bred in-house. Primers used for genotyping were Casp8 (5′-GGATGTCCAGGAAAAGATTTGTGTC-3′, 5′-CCTTCCTGAGTACTGTCACCTGT-3′) and RIPK3 (5′-CGCTTTAGAAGCCTTCAGGTTGAC-3′, 5′-GCAGGCTCTGGTGACAAGATTCATGG-3′, 5′-CCA-GAGGCCACTTGTGTAGCG-3′) (Integrated DNA Technologies).

Pup sexing was based on differences in anogenital pigmentation: male pups exhibit a pigmented spot that eventually develops into the scrotum, while female pups lack this marker^87^.

### Brain tissue processing and single-cell dissociation

P4 WT, RIPK3 KO, and DKO mice were decapitated, and their brains were dissected into four regions: telencephalon, diencephalon, mesencephalon, and rhombencephalon. Single-cell separation was achieved using a modified version of Volovitz and colleagues’ method^88^. Brain tissue was enzymatically digested in a solution containing dispase II (Sigma-Aldrich, D4693), collagenase type IV (Worthington, LS004186), DNAse-I (Sigma-Aldrich, 11284932001), and hyaluronidase (Sigma-Aldrich, H3884). After mechanical dissociation using a P1000 pipette and 30-minute incubation at 37°C, the cell suspension was filtered through a 45-μm mesh, centrifuged at 300 x g for 5 minutes at 4°C, and resuspended in DPBS (DPBS; Thermo Fisher Scientific, 14190) for subsequent Cisplatin staining.

### Cisplatin staining, fixation and cell counting

Cells were incubated with 10 µM Cisplatin (Sigma Aldrich, P4394) for 30 seconds. The reaction was quenched and diluted 1:10 with DPBS containing 0.5% bovine serum albumin (BSA; Sigma-Aldrich, A9418) and immediately centrifuged at 300 x g for 3 minutes at 4°C. After centrifugation and resuspension, cells were fixed with 1.6% paraformaldehyde (PFA; Electron Microscopy Services, CAS 30525-89-4) for 10 minutes. Following a DPBS wash, cells were prepared in cell staining medium (CSM; 0.5% BSA, 0.02% NaN_3_ in PBS). A 100 μL aliquot was reserved for flow cytometry quality control, and the remaining sample was stored at-80°C. Dissociation quality was first confirmed by microscopic inspection using an EVOS AMF4300 microscope and later through flow cytometry. Samples with predominantly single cells and minimal debris were counted using a Bio-Rad TC20 Automated Cell Counter.

### Sample barcoding, staining, and intercalation for mass cytometry

Telencephalon samples from P4 RIPK3 KO mice, DKO mice, and WT mice were processed for mass cytometry using an established protocol^33^. Samples were barcoded, pooled for antibody staining, and intercalated. The surface and intracellular antibodies are listed in Supplementary Table 1. The analysis was conducted on a CYTOF XT System (Fluidigm Corporation) by the University of Virginia Flow Cytometry Core.

### Normalization, debarcoding, gating, and compensation

Raw FCS data files were standardized through bead normalization and debarcoded^89,90^. Single-cell isolation was performed via cleanup gating using Cytobank (https://community.cytobank.org/cytobank/experiments/116996) (Supplementary Fig. S7C-E). To minimize signal spillover between channels, a manual compensation matrix was applied^91^, adapted from Chevrier et al. The resulting high-quality, single-cell data for each sample were exported as individual.fcs files for subsequent analysis. To ensure reproducibility and transparency, all raw data files are publicly available through Flow Repository (https://flowrepository.org/) [ID: <u>FR-FCM-Z934</u>].

### Batch correction

To mitigate batch effects and ensure consistent antibody signals across barcode sets, batch correction was performed^92^. A universal sample containing cells from all three genotypes was used to standardize signals. Antibodies producing Gaussian distributions with mean signal variance >1% were corrected at the 50th percentile (e.g., PDGFRa). CD133, Ki67, and GFAP were corrected at the 95th percentile. Other markers with low variance (<1%) were not batch corrected.

### Clustering of high-dimensional data

Cells were clustered using 37 expression markers, with populations identified based on established CNS marker profiles^33^. Then data were dimensionally reduced using uniform manifold approximation and projection (UMAP) for visualization ^55,93^, with parameters: nearest neighbors = 15, metric = Euclidean, local connectivity = 1, components = 2, epochs = 1000.

### Tissue preparation and immunohistochemistry

P4 mouse brains were fixed (4% PFA, 48 hours), dehydrated (30% sucrose, 72 hours), frozen (-80°C), embedded in OCT, and cryosectioned (30-μm).

Mounted sections were washed with PBS, blocked (2% normal donkey serum, 1%BSA, 0.1% Triton, 0.05% Tween-20 in PBS, 1h), and incubated with primary antibodies overnight at 4°C: PECAM (PE anti-mouse CD31, rat, 1:300, BioLegend, catalog 160203), NeuN (NeuN, rabbit, 1:1,000, Abcam, catalog ab177487) and Tbr2 (EOMES, rat, 1:300, Invitrogen, catalog 14-4875-82). After PBST (PBS + 0.05% Tween 20) washes, sections were incubated with Alexa Fluor secondary antibodies (1:500; Thermo Fisher Scientific; 2h), washed with PBS, and mounted with Fluoromount-G (SouthernBiotech, catalog 0100-01).

### RNAscope

Brain sections on charged slides were air-dried (36h), treated with H_2_O_2_ (RNAscope H2O2 and Protease Reagents Kit, Advanced Cell Diagnostics, catalog 322381, 10min), and incubated with protease IV (RNAscope H2O2 and Protease Reagents Kit, Advanced Cell Diagnostics, catalog 322381, 15min, 40°C). After washing, tissue was incubated in Probe Master Mix for 2 hours at 40°C (Probe1: Mm-RipK3, catalog 462541; Probe2: Mm-Casp8-C2, catalog 468971-C2; Advanced Cell Diagnostics). Samples were then amplified using AMP1-3 and visualized with TSA Cyanine 5 and TSA Cyanine 3 staining.

### Image analysis

Images were taken using a Zeiss 980 confocal laser-scanning microscope at ×20/0.8 magnification. All confocal microscopy images were processed using the “Fiji is Just ImageJ” application and set to maximum intensity projection.

For Tbr2^+^ cell number quantification, images were separated into individual channels, and the Tbr2 channel was isolated for further analysis. Brightness and contrast adjustments were made to maximize positive signal intensity and minimize background intensity. The entire image was then analyzed using the “QuPath” application. The “Polygon” tool was used to outline the region of interest (ROI) and the “Cell detection” tool to identify Tbr2^+^ cells. Manual checks were performed to ensure that all positive cells were detected accurately. In total,2 images from each section with 3 sections per animal were included in the analysis.

For PECAM^+^ area quantification, images were again separated into individual channels, and the PECAM channel was isolated for further analysis. Image brightness and contrast were adjusted to optimize signal detection. The Labkit Pixel Classification plugin was used to classify the entire image, with representative foreground and background pen markings adjusted manually to improve positive signal detection accuracy. The segmentation result was reimported to ImageJ and thresholded to separate “foreground” and “background” pixel classes. Each section image was analyzed independently. The Analyze > Measure function was used to quantify the %Area positive for PECAM. In total, 3 images from each section with 3 sections per animal were included in the analysis.

### Statistical Analysis

For time course experiments, developmental trajectories were fitted using locally weighted regression (LOESS) with 95% confidence intervals. R² values were calculated as 1 - (SSresidual/SStotal), where SSresidual represents the sum of squared differences between observed and LOESS-predicted values, and SStotal represents the sum of squared differences between observed values and their mean.

Statistical comparisons among WT, RIPK3 and DKO were primarily performed using one-way ANOVA followed by Tukey’s post-hoc test for multiple comparisons For direct comparisons between two groups in heatmap, Student’s t-test was applied. Statistical significance levels were defined as: *p < 0.05, **p < 0.01, ***p < 0.001, with non-significant differences denoted as “ns”.

All statistical analyses were performed using R (4.4.1) and Python (3.8) with standard statistical packages. Graphs were generated using ggplot2 in R.

## Acknowledgments

Research reported in this publication was supported by the National Institute of Neurological Disorders and Stroke of the National Institutes of Health grant R01NS111220 (awarded to E.R.Z. and C.D.D.), and the Owens Family Foundation (awarded to C.D.D. and J.R.L.). The content is solely the responsibility of the authors and does not necessarily represent the official views of the National Institutes of Health. Further support was provided by the 3Cavaliers Pilot Research Program to C.D.D. and E.R.Z. We thank all members of the Deppmann lab for helpful discussions, Stefani Mancuso for editorial assistance, and Anthony Spano for technical expertise. We also acknowledge the University of Virginia Flow Cytometry Core for technical support with the CYTOF XT System. We appreciate the contributions of the University of Virginia Research Computing for providing computational resources and technical support that have contributed to the results reported in this publication.

## Author Contributions

J.S., C.D.D., and E.R.Z. designed all experiments. J.S. conducted all experiments with assistance from W.L., A.S., and T.S. A.V.D. performed antibody conjugations and validated antibodies by conducting titration experiments on positive and negative control cell samples. J.S., W.L., and A.V.D. performed mass cytometry measurements. A.S. and T.S. performed immunohistochemistry (IHC) and RNAscope in situ hybridization (ISH) measurements. J.S., A.S., and T.S. conducted microscopy. J.S. and E.R.Z. wrote scripts for data analysis. J.S., A.S., T.S. and E.R.Z. performed data analysis. E.R.Z. and C.D.D. conceived and supervised all aspects of the project. J.S., A.S., T.S., and E.R.Z. prepared figures. J.S., E.R.Z., and C.D.D. wrote the manuscript with input from all authors.

## Declaration of Generative AI and AI-Assisted Technologies in the Writing Process

During the preparation of this work, the author(s) used [ChatGPT] and [Claude from Anthropic] in order to assist with drafting and refining the manuscript text. After using these tools or services, the author(s) reviewed and edited the content as needed and take(s) full responsibility for the content of the publication.

---

Supplementary Fig. S1 Cell Death, Proliferation, and Abundance in the Developing Telencephalon.

(A) Representative flow cytometry plots showing distinct cell populations based on CC3 and Cisplatin expression patterns during development: CC3^+^Cisplatin^-^ (early apoptosis), CC3^-^ Cisplatin^+^ (non-apoptotic death), and CC3^+^Cisplatin^+^ (late-stage apoptosis or mixed death mechanisms).

(B) Violin plots showing marker expression distribution across 23 cell clusters (rows) for 37 protein markers (columns).

(C) Analysis of neuronal cluster dynamics during development. Top: UMAP visualizations highlighting individual clusters categorized into immature and mature neuronal populations. Bottom: Temporal profiles from E13 to P4 showing cluster abundance (gray), percentage of CC3^+^ cells (orange), Cisplatin^+^ cells (green), and Ki67^+^ cells (blue). Points represent individual replicates, and solid lines show Loess curve fitting of the data.

Supplementary Fig. S2 Analysis of Nonneuronal Subpopulations Dynamics during Development.

Top: UMAP visualizations highlighting individual clusters categorized into immature and mature nonneuronal populations. Bottom: Temporal profiles from E13 to P4 showing cluster abundance (gray), percentage of CC3^+^ cells (orange), Cisplatin^+^ cells (green), and Ki67^+^ cells (blue). Points represent individual replicates, and solid lines show Loess curve fitting of the data.

Supplementary Fig. S3 Antibody Testing of Cleaved Caspase-8 (CC8) by IHC during Telencephalic Development.

Representative immunofluorescence images of mouse brain sections from E13.5 to P0. Sections were stained with DAPI (blue), cleaved Caspase-3 (CC3) (red, Cell Signaling #9661S), and two different CC8 antibodies: Cleaved Caspase-8 (Asp387) (D5B2) XP® Rabbit mAb, Cell Signaling #8592S and Cleaved Caspase-8 (Asp384) (11G10) Mouse mAb, Cell Signaling #9748S. Multiple developmental timepoints were examined to assess CC8 antibody specificity and optimize staining conditions. Scale bar: 50 μm.

Supplementary Fig. S4 Antibody Testing of pRIPK3 and pMLKL by IHC Using WT, RIPK3 KO and DKO Brain Sections.

(A) Representative immunofluorescence images of mouse brain sections from WT and DKO mice. Sections were stained with DAPI (blue) and pMLKL (Phospho-MLKL (Ser345) (D6E3G) Rabbit mAb, Cell Signaling #37333S, red).

(B) Representative immunofluorescence images of mouse brain sections from WT, RIPK3 KO and DKO mice. Sections were stained with DAPI (blue) and pRIPK3 (Phospho-RIP3 (Thr231/Ser232) Antibody (Mouse Specific), Cell Signaling #57220S, red). Scale bar: 50 μm.

Supplementary Fig. S5 Representative Images of RNAscope In Situ Hybridization for Caspase-8 (Magenta) and Ripk3 (Yellow) mRNA in 6 Month Mouse Cortex.

Images were captured at both 20× and 40× magnification to show both broad distribution patterns and cellular-level expression details. Scale bar: 50 μm.

Supplementary Fig. S6 Sex-specific analysis in P4 telencephalon.

(A) Quantification of total cell numbers in the telencephalon of P4 male (M) and female (F) mice across genotypes (WT, RIPK3 KO, and DKO; n = 6 per group).

(B) Quantification of CC3^+^, Cisplatin^+^, and Ki67^+^ cells% in the telencephalon of P4 male (M) and female (F) mice across genotypes (WT, RIPK3 KO, and DKO; n = 6 per group). Data presented as box plots where the middle line represents the median, the box represents the interquartile range (IQR), and the whiskers extend to 1.5 times the IQR. Individual data points are shown as dots. Statistical differences between groups are denoted by asterisks.

Supplementary Table 1. Mass Cytometry Antibody Panel.

Comprehensive list of antibodies used in mass cytometry analysis showing metal isotope labels (Metal), antibody name (Antibody/Reagent), and full protein names or descriptions (Full Name(s)), vendor information (Vendor), catalog numbers (Catalog No.), clone identifiers (Clone), working concentrations (concentration).

Supplementary Fig. S7. Pre-processing Workflow for Mass Cytometry Data

(A) Raw data normalization using calibration beads, showing signal intensities before and after normalization.

(B) Debarcoding analysis of normalized data across three barcode sets, with event count distributions (top) and yield percentages (bottom) for each sample.

(C-E) Sequential gating strategy for single-cell isolation: barcode stringency (87.98%), event-centered mass cytometer isolation (94.16%), and bead removal (98.64%). Gates shown in red with percentages indicating cell yield.

(F) Signal compensation matrix used to correct for metal isotope spillover between channels.

(G) Comparison of mean signal intensities before (blue) and after (yellow) batch correction for CD133, Ki67, and GFAP. Only markers with mean signal variance > 0.01 underwent batch correction; all other markers showed minimal variance and remained uncorrected.

Supplementary Fig. S8 Cell Type-specific Markers Used for Mass Cytometry Analysis.

Overview of protein markers used to define distinct cell populations: neural stem cells and progenitors (blue, top left), general neuronal identity (blue, bottom left), neuronal lineage (blue, top middle), glial lineages (yellow, bottom middle), and nonneuronal cell types including microglia, oligodendrocytes, astrocytes, and endothelial cells (orange, right).

Supplementary Fig. S9 Refined Clustering Analysis of Mass Cytometry Data.

(A) UMAP visualization showing 89 distinct cell clusters identified by secondary Leiden clustering analysis. Clusters are color-coded and numbered (1-89) as shown in the legend.

(B) Violin plots depicting marker expression distribution across all 89 clusters (rows) for 37 protein markers (columns).

Supplementary Fig. S10 Genotype-Dependent Abundance Differences Across Initial Clusters in the Developing Telencephalon.

Heatmap showing marker gene expression across 30 identified cell clusters (rows) from the initial cluster with corresponding p-values comparing relative abundance between genotypes (DKO vs. WT, RIPK3 KO vs. WT and DKO vs. RIPK3 KO). The main heatmap shows gene expression levels where yellow indicates higher expression and blue indicates lower expression (scale shown on right). The adjacent p-value heatmap (3 right columns) displays statistical significance of abundance differences between genotypes (DKO vs. WT, RIPK3 KO vs. WT and DKO vs. RIPK3 KO), where red indicates higher relative abundance and blue indicates lower relative abundance (scale shown on right). For all quantifications, statistical differences were calculated using Student’s t-test.

