## Supplemental Table 1 for "Extrinsic Apoptosis and Necroptosis in Telencephalic Development: A Single-Cell Mass Cytometry Study"

| Metal | Antibody/Reagent | Full Name(s) | Vendor | Catalog No. | Clone | Concentration |
|---|---|---|---|---|---|---|
| Y89 | TUJ1 | Beta 3-tubulin, Tuj1 | Gift (A. Spano) | - | Tuj1 | 500 ng/mL |
| In113 | Olig2 | Oligodendrocyte transcription factor 2 | Millipore | MABN50 | 211F1.1 | 3000 ng/mL |
| Pr141 | A2B5 | - | Biolegend | 150702 | 105/A2B5 | 50 ng/mL |
| Nd143 | PECAM1 | PECAM-1, Endothelial cell adhesion molecule | Biolegend | 102425 | 390 | 50 ng/mL |
| Nd144 | Nestin | - | R&D Systems | MAB2736 | 307501 | 10 ng/mL |
| Nd145 | Sox1 | SRY-box transcription factor 1 | R&D Systems | AF3369 | Polyclonal | 80 ng/mL |
| Nd146 | Tbr2 | T-box brain gene 2, EOMES, eomesdermin | Thermo Fisher | 14-4875-82 | Dan11mag | 4000 ng/mL |
| Sm147 | PDGFRa | Platelet-derived growth factor alpha | Biolegend | 135902 | APA5 | 100 ng/mL |
| Nd148 | CD133 | Prominin 1 | Biolegend | 141202 | 315-2C11 | 80 ng/mL |
| Sm149 | Islet1 | ISL LIM homeobox 1 | Novus | NBP2-14999 | Polyclonal | 250 ng/mL |
| Nd150 | RBFOX3/NeuN | Neuronal Nuclei | Novus | NBP1-92693 | 1B7 | 200 ng/mL |
| Eu151 | Sox10 | Sex-determining region Y box 10 | Gift (S. Kucenas) | - | Monoclonal | 500 ng/mL |
| Sm152 | Ki67 | MKI67, Marker of proliferation Ki-67 | BD Biosciences | 550609 | B56 | 35 ng/mL |
| Eu153 | OligoO4 | Oligodendrocyte marker 04 | R&D Systems | MAB1326 | O4 | 250 ng/mL |
| Sm154 | Pax6 | Paired box 6 | BD Biosciences | 561462 | O18-1330 | 15 ng/mL |
| Gd155 | PSA-NCAM | Polysialylated-neural cell adhesion molecule | eBioscience | 14-9118-82 | 12E3 | 150 ng/mL |
| Gd156 | GLAST | Excitatory amino acid transporter 1,Glutamate asparate transporter | Novus | NB100-1869 | Polyclonal | 15 ng/mL |
| Gd157 | SSEA-1 | CD15, Stage-specific embryonic antigen | eBioscience | 14-8813-80 | MC-480 | 8 ng/mL |
| Gd158 | CD11b | Integrin aM, Mac-1 | Biolegend | 101249 | M1/70 | 15 ng/mL |
| Tb159 | CD24 | Heat stable antigen | BD Biosciences | 557436 | m1/69 | 10 ng/mL |
| Gd160 | Sox2 | Sex-determining region Y bax 2 | R&D Systems | MAB2018 | 245610 | 3000 ng/mL |
| Dy161 | N-Cadherin | N-cadherin, CD325 | Biolegend | 844702 | 13A9 | 30 ng/mL |
| Dy162 | GAD65 | Glutamic acid decarboxylase 65-kD, glutamate decarboxylase 2 | Biolegend | 844502 | N-CAD65 | 600 ng/mL |
| Dy163 | Dcx | Doublecortin | Thermo Fisher | 481200 | Polyclonal | 500 ng/mL |
| Dy164 | MAP2 | Microtubule-associated protein 2 | Novus | NBP2-25156 | 4H5 | 200 ng/mL |
| Ho165 | ALDH1A1 | Aldehyde dehydrogenase 1A1 | R&D Systems | AF5869 | Polyclonal | 80 ng/mL |
| Er166 | Ly-6C | Lymphocyte antigen 6 complex, locus C | Biolegend | 128002 | HK1.4 | 10 ng/mL |
| Er167 | CD45 | Protein tyrosine phosphatase receptor type C | Fluidigm | 3089005B | 30-F11 | 10 ng/mL |
| Er168 | Ctip2 | COUP-TF-interacting protein 2, Bcl11b | Abcam | ab18465 | 25B6 | 200 ng/mL |
| Tm169 | GFAP | Glial fibrillary acidic protein | BD Biosciences | 556330 | 102 | 7 ng/mL |
| Er170 | Cux1 | Cut-like homeobox 1 | Abcam | ab54583 | 2A10 | 10 ng/mL |
| Yb171 | Tbr1 | T-box brain gene 1 | Abcam | ab31940 | Polyclonal | 1000 ng/mL |
| Yb172 | BLBP | Brain lipid-binding protein, fatty acid binding protein 7, FABP7 | Gift (C. Birchmeier | - | Polyclonal | 1000 ng/mL |
| Yb173 | cl-Casp3 | cleaved Caspase-3 | BD Biosciences | 559565 | C92-605 | 2000 ng/mL |
| Yb174 | TrkB | Neurotrophic tyrosine kinase receptor type 2 | Thermo Fisher | AF1494 | Polyclonal | 300 ng/mL |
| Lu175 | F4/80 | EMR1, Ly-71 | Biolegend | 123101 | BM8 | 5 ng/mL |
| Yb176 | p75NTR | P75 neurotrophic receptor, TNF receptor 16 | R&D Systems | AF1157 | Polyclonal | 150 ng/mL |
| Ir191/193 | Intercalator | Cell-ID Intercalator-Ir | Fluidigm | 201192A | - | 1:5000 dilution |
| Pt194/195/198 | Cisplatin | Cisplatin | Sigma Aldrich | P4394 | - | 5 µM |
